# Functional effects of deforestation on stream ecosystems in Madagascar based on growth and production of mayflies (Ephemeroptera)

**DOI:** 10.1101/2025.01.17.633570

**Authors:** Ranalison Oliarinony, Michel Sartori

**Author notes:** **Correspondence:** Michel Sartori, Naturéum, Department of zoology, Palais de Rumine, Place Riponne 6, CH-1005 Lausanne, Switzerland.

## Abstract

The effects of deforestation on species richness and abundance of tropical freshwater organisms are increasingly well documented, but corresponding effects on functional properties of ecosystems, such as productivity and growth, are largely unknown. Here we investigated the biomass production and growth of eight species of mayflies (Ephemeroptera) in streams of eastern Madagascar. We compared three study sites in primary tropical forest with three in degraded open canopy area (*savoka*). All species were asynchronous and aseasonal. Seven species were multivoltine (3-4 generations per year) and one was semi-voltine. Biomass production of the most abundant species was 4-5 times greater in *savoka* than in forest, and standing biomass: productivity (P/B) was always higher in *savoka*. This production shift was mainly caused by the two *Xyrodromeus* species, both of which feed on periphytic algae that are more abundant in an open canopy. Surprisingly, *Xyrodromeus* females emerged more rapidly and were larger in size in *savoka* than in forest. Our results clearly indicate that deforestation in tropical Madagascar leads not only to a shift in mayfly community composition, but perhaps more importantly to functional changes in the ecosystem, namely standing biomass, annual production, and female size at maturity.

## Introduction

Tropical deforestation results from logging, agriculture, (Flamenco-Sandoval et al. 2007), mining, (Akpalu and Parks 2007) and road construction (Goosem 2007). Beyond the immediate effects to forest ecosystems, deforestation has severe consequences for the biodiversity of streams and rivers (Dudgeon 2000, 2003, Iwata et al. 2003). Despite representing < 0.01% of the world’s water and 0.8% of the earth’s surface, freshwaters harbour ca. 10% of known animal biodiversity (Dudgeon et al. 2006, Balian et al. 2008). Madagascar is a well known biodiversity hotspot (Goodman and Benstead 2005) but ongoing deforestation has led to a 40% decrease in primary forest area between 1950 and 2000 (Harper et al. 2007). The consequences are increasingly well studied for terrestrial communities (Irwin 2008), but poorly known for freshwater fish and invertebrates (Benstead et al. 2003). Mayflies (Ephemeroptera) play an important role in freshwater ecosystems as primary consumers (Malmqvist 2002), and their biodiversity on Madagascar is unique, with all but one species endemic to the island (Elouard et al. 2003, Barber-James et al. 2008).

While quantitative data on the effects of deforestation on species richness and abundance in Afrotropical freshwaters are scarce, functional data such as biomass production and growth rates are unavailable (Jacobsen et al. 2008). There are relatively few reports of biomass production in the tropics (Boulton et al. 2008) and those available for mayflies represent only *ca*. a dozen species.

We studied the relative abundance, life cycle, and biomass production of eight endemic mayfly species in forested and deforested areas of eastern Madagascar. From the few data available in other tropical areas, we predicted aseasonality and asynchronous life cycles in Afrotropical mayflies. We also tested whether life cycles were shorted in deforested *savoka* compared with forest as expected based on early works (e.g.Sweeney and Vannote 1978).

## Methods

### Study sites

We studied six second- or third-order streams in the Rianila watershed (7820km^2^, Fig. 1, Table 1). All were located in the Eastern Domain, once entirely covered with rainforest, and now heavily altered by human activities. Mean annual rainfall is ca 1790mm and mean air temperature 18°C. (Chaperon et al. 1993). The area is characterised by a “dry” season from April until October when daily rain falls or showers are common, and a rainy season from November to March.

**Fig. 1:**
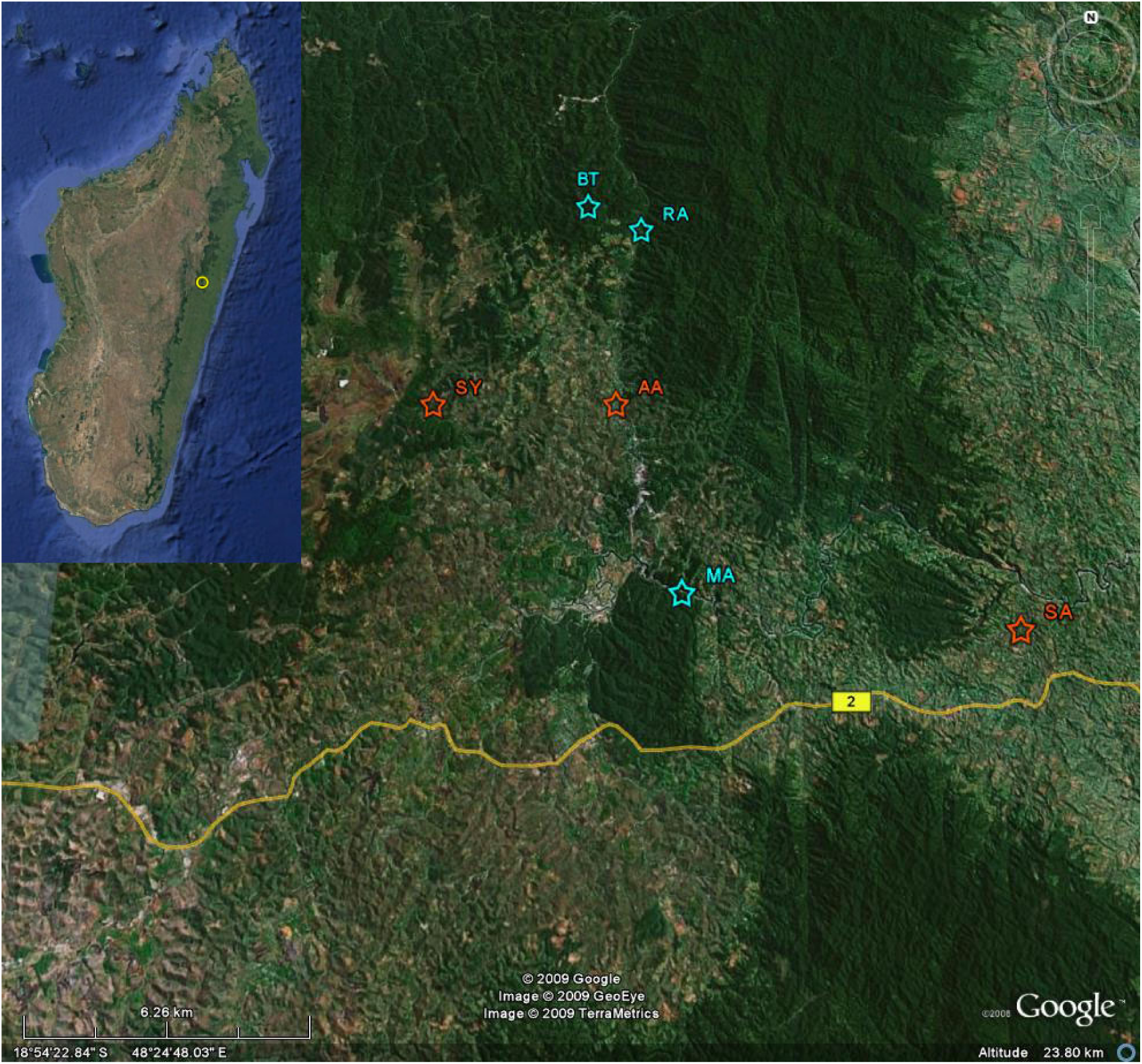
Location of the study sites on Google Earth © representation (yellow ring in inset). Sites label according to their abbreviation in table 1; blue stars and orange stars indicate position in forest (dark green background) and in savoka (light green background) respectively. Yellow line indicates position of National Road #2 from Antananarivo to Toamasina.

**Table 1:**
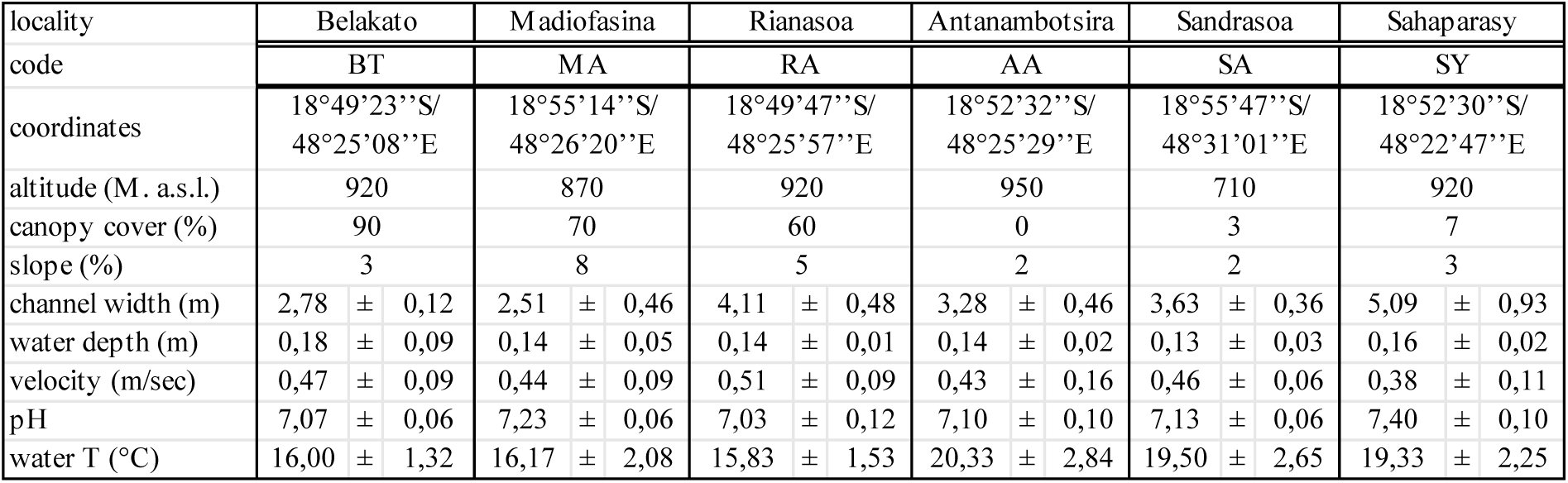
Environmental characteristics of the study sites.

Three sites (BT, MA, RA) were located in primary forest of the Mantady National Park, and three (AA, SA, SY) were located in degraded, open canopy areas composed of *Tavy* (slash-and-burn agriculture, mainly for rice) and *Savoka* (fallow fields). Deforestation typically leads to a replacement of native species by exotics and favours grasses over woody species, creating treeless landscapes that are of minimal productive and ecological value (Styger et al. 2007).

### Sampling

Samples were collected each month from June 2001 until June 2002, with the exception of November 2001 (bimonthly samplings) and March 2002 (no samplings due to heavy rains at the end of the wet season). We took three samples at each site using a Surber net (0.1 m^2^, mesh 250 µm) and preserved samples in the field with 70% ethanol.

### Laboratory methods

We focused on eight mayfly species that were abundant and could be identified even at early (small) larval stages (Table 2). These belonged to five families and seven genera. Head width of all specimens was measured to the nearest 0.1 mm using a stereomicroscope and an eyepiece micrometer. Mature nymphs (recognizable by their black wing pads) were sexed and measurements used to investigate sizes at emergence. Density was measured as the mean number of individuals m^-2^ (*n* = 3) and biomass was derived from mean density and length-mass relationships reported by Benke et al. (1999). Results are expressed in mg dry weight (DW) m^-2^. For *Proboscidoplocia cf. vayssierei* and *Elassoneuria cf. insulicola*, no useful data were available in the literature, and we derived new equations (Table 2) using specimens of all size classes (*n*=109 and *n*=102 respectively) dried for 24 h at 60°C, cooled in dessicator, and weighed to the nearest 0.01 mg on an analytical balance.

**Table 2:**
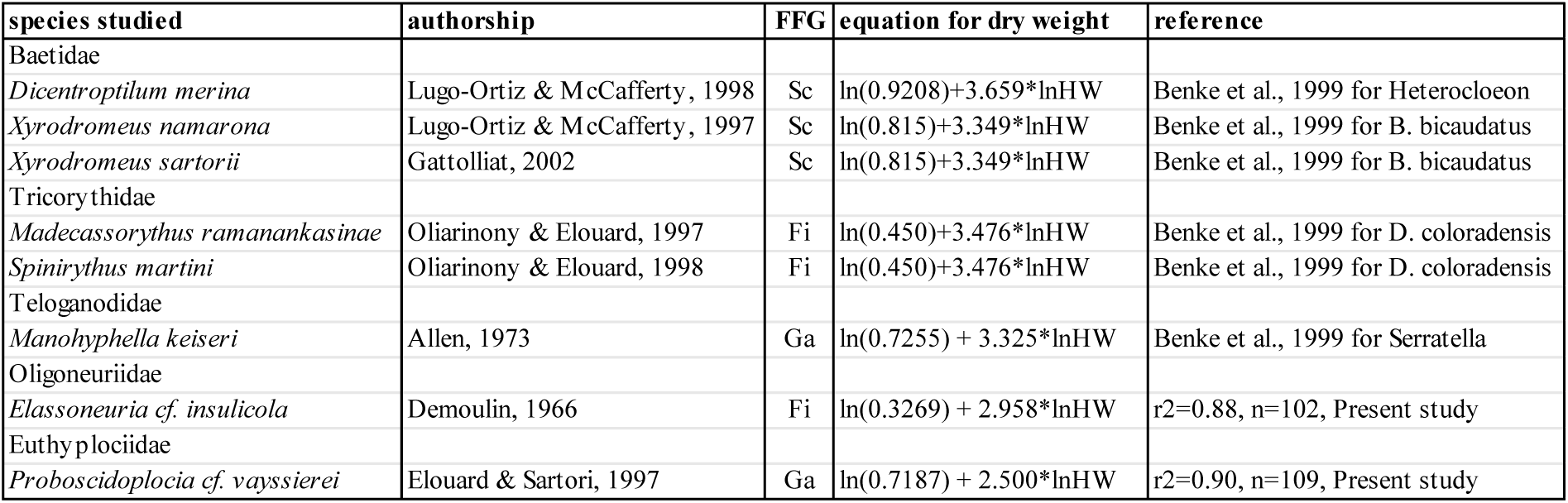
List of studied species, arranged by family, their functional feeding group (FFG) and references of the equation used to calculate their dry weight; Sc = scraper, Ga = collector-gatherer, Fi = filter; HW = head width (in mm).

All species investigated had aseasonal and asynchronous life cycles (see Results). Production was therefore calculated using the instantaneous growth method and the size-frequency method (Benke 1993, Morin and Dumont 1994, Marchant and Yule 1996). Equations used were:

### Instantaneous growth method

Log g = −2.07 + 0.038 T-0.14log DW, where g is the daily growth rate (day^-1^), T is the temperature of the water (in °C), and DW the dry weight (in mg). For *Proboscidoplocia cf. vayssierei* the peculiar morphology led us to use the regression equation proposed for Plecoptera rather than Ephemeroptera (Morin and Dumont 1994), so the equation was corrected to: log g = −1.9 + T-0.28log DW.

### Size-frequency method

LogP = log B+0.716+0.030 T-0.382log Wm where P is the production (mg DW m^-2^ y^-1^), B is the biomass (mg DW m^-2^), and Wm is the maximum dry weight of the species. As for the instantaneous growth method, we used the regression equation proposed for Plecoptera for *Proboscidoplocia cf. vayssierei*: logP = log B+0.592+0.015 T-0.16log Wm.

Production for each cohort (CP) was calculated and larval life span or cohort production interval (CPI) was estimated according to Marchant and Yule (1996). Degree day demand (DDD) and P/B ratio were also calculated. For comparative purposes, data from the literature reported as Ash Free Dry Weight (AFDW) were converted into DW using DW = 1.11 AFDW (Benke 1993).

Functional feeding group was assigned to each of the eight species, based on morphological characteristics and available literature; Oligoneuriidae (brush-legged mayflies), and Tricorythidae (Barber-James and Lugo-Ortiz 2003) were assigned to the filter-feeding guild, Euthyplociidae (Fenoglio et al. 2008) and Teloganodidae (McCafferty and Benstead 2002) were considered to be collector-gatherers, and the Baetidae *Xyrodromeus* (Gattolliat 2002) and *Dicentroptilum* (J.-L. Gattolliat comm. pers.) were assigned to the algal scraping guild.

## Results

### Density

Among the 8 species studied, the 3 Baetidae were the most abundant and were the only species abundant enough to be studied in detail in all localities. *E. cf. insulicola* was studied in one forest and one *savoka* locality, whereas *M. ramanankasinae* and *M. kaiseri* were abundant only in forest and *P. cf. vayssierei* and *S. martini* only in *savoka* (Table 3).

**Table 3:** Mean densities per square meter (± SD) of the mayfly species in the study sites; abbreviations as in. table 1.

|  | BT | MA | RA | AA | SA | SY |
| --- | --- | --- | --- | --- | --- | --- |
| *D. merina* | 9,72 ± 12,18 | 11,39 ± 12,02 | 16,11 ± 22,64 | 54,17 ± 49,01 | 44,17 ± 26,60 | 14,44 ± 12,34 |
| *X. namarona* | 31,11 ± 25,52 | 74,17 ± 58,38 | 59,72 ± 95,86 | 162,78 ± 99,23 | 190,56 ± 149,98 | 62,22 ± 70,29 |
| *X. sartorii* | 56,11 ± 25,66 | 110,56 ± 82,13 | 16,11 ± 18,95 | 80,83 ± 72,62 | 336,11 ± 284,38 | 156,94 ± 149,56 |
| *M. ramanankasinae* | | | 16,01 ± 9,82 | | | |
| *S. martini* | | | | | 30,83 ± 42,76 | |
| *E. cf. insulicola* | | | 20,83 ± 13,19 | | 19,72 ± 32,80 | |
| *P. cf. vayssierei* | | | | | 35,00 ± 26,15 | |
| *M. kaiseri* | | 12,50 ± 17,93 | 10,56 ± 11,18 | | | |

Densities were highly variable among sampling dates (high standard deviation) and no correlation could be drawn between seasons. *Savoka* sites generally had higher mean densities than forested sites (up to 3 times for the Baetidae species), but differences were not significant, largely due to the low density at deforested site SY (Fig. 3A).

### Life cycle

All species had an asynchronous and aseasonal life cycle, with almost constant recruitment and multiple size classes present throughout the year (Fig. 2).

**Fig. 2:**
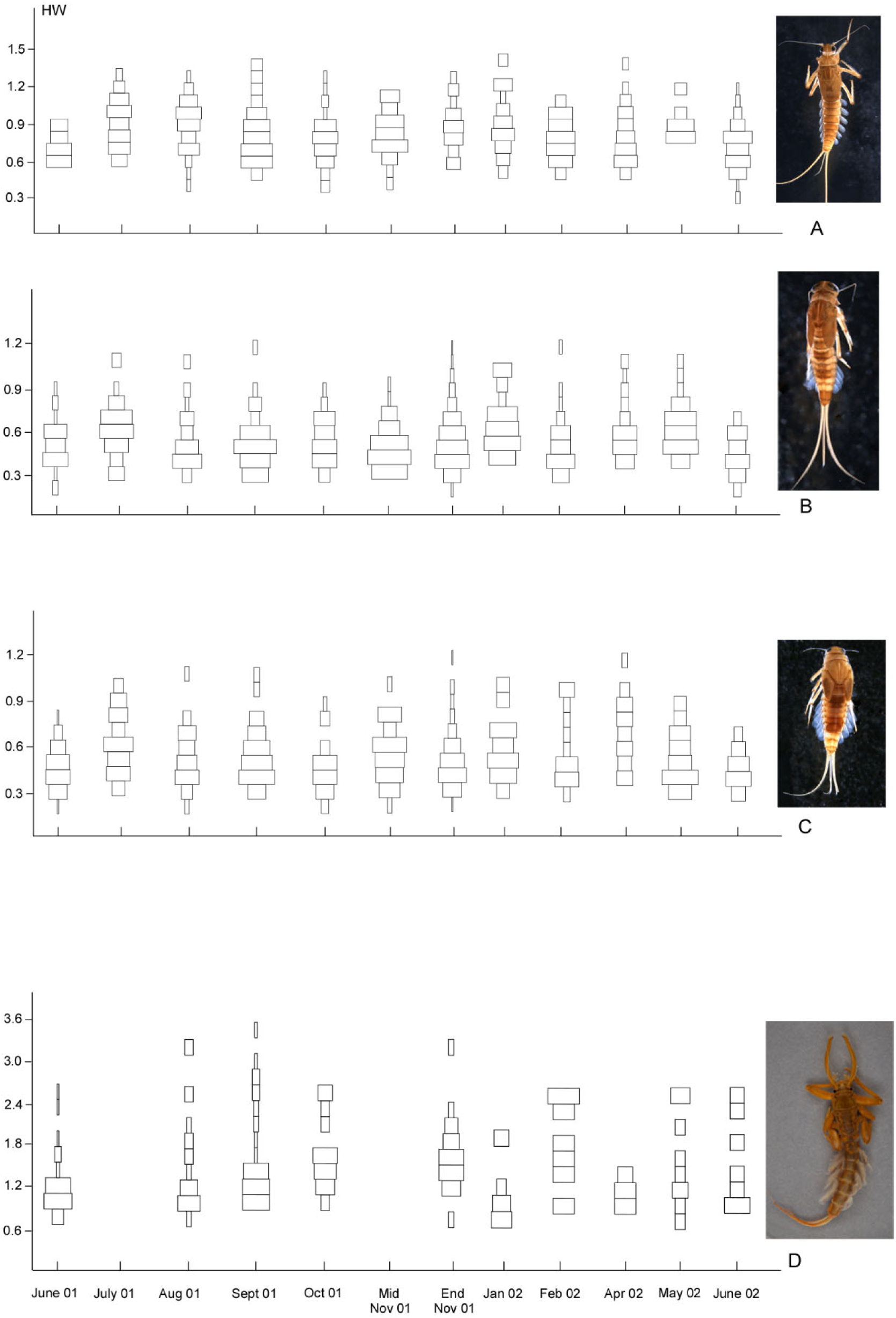
Examples of life cycle graphs obtained in forested or savoka areas. A: *D. merina* in a savoka site; B: *X. sartorii* in a forest site, C: *X. namarona* in a forest site, D: *P. cf. vayssierei* in a savoka site. HW: head width.

CPI showed a development time of 3-4 months for the 3 Baetidae species, but reached 12-18 months, depending of the method used, for *P. cf. vayssierei* (Table 4). *D. merina* exhibited no significant differences in CPI between savoka and forest sites, in contrast to the two *Xyrodromeus* species where the life cycle in *savoka* was *ca*. 20 d shorter than in forest (*p* = 0.017 and *p* = 0.0099 for *X. namarona* and *X. sartorii* respectively). Degree-day demands were comparable among all studied populations, ranging from ca 1600 to 2000, with the notable exception of *P. cf. vayssierei* whose long life cycle requires more than 10000 DD to be completed.

**Table 4:** Mean biomass and production of the investigated species; F: forest; S: savoka, biomass expressed in mg DW m^-2^; production calculated by two methods and expressed in mg DW m^-2^ y^-1^; CPI: cohort production interval (in days); DDD: degree-day demand.

|  |  |  | Instantaneous growth method |  |  |  | Size-frequency method |  |  |  |
| --- | --- | --- | --- | --- | --- | --- | --- | --- | --- | --- |
|  |  | biomass | production | P/B | CPI | DDD | production | P/B | CPI | DDD |
| *D. merina* |  |  |  |  |  |  |  |  |  |  |
| BT | F | 3,87 | 53,09 | 13,72 | 104 | 1661 | 43,46 | 11,23 | 127 | 2029 |
| MA | F | 3,16 | 46,14 | 14,58 | 107 | 1729 | 45,35 | 14,33 | 109 | 1759 |
| RA | F | 9,53 | 121,81 | 12,78 | 99 | 1574 | 118,94 | 12,48 | 102 | 1612 |
| AA | S | 24,37 | 474,28 | 19,46 | 92 | 1875 | 339,19 | 13,92 | 129 | 2621 |
| SA | S | 20,79 | 376,99 | 18,14 | 99 | 1933 | 273,24 | 13,14 | 137 | 2667 |
| SY | S | 8,12 | 141,96 | 17,47 | 82 | 1584 | 105,55 | 12,99 | 110 | 2130 |
| *X. namarona* |  |  |  |  |  |  |  |  |  |  |
| BT | F | 2,87 | 45,30 | 15,77 | 111 | 1776 | 48,67 | 16,94 | 103 | 1653 |
| MA | F | 6,86 | 113,33 | 16,51 | 114 | 1842 | 135,21 | 19,69 | 95 | 1544 |
| RA | F | 6,36 | 98,75 | 15,52 | 126 | 1992 | 94,55 | 14,86 | 131 | 2081 |
| AA | S | 14,94 | 363,45 | 24,32 | 89 | 1816 | 341,42 | 22,85 | 95 | 1933 |
| SA | S | 18,53 | 408,37 | 22,03 | 103 | 2002 | 354,93 | 19,15 | 118 | 2303 |
| SY | S | 8,61 | 181,45 | 21,08 | 91 | 1759 | 162,91 | 18,93 | 101 | 1959 |
| *X. sartorii* |  |  |  |  |  |  |  |  |  |  |
| BT | F | 6,61 | 102,27 | 15,48 | 121 | 1937 | 99,38 | 15,04 | 125 | 1994 |
| MA | F | 15,52 | 243,33 | 15,67 | 118 | 1909 | 236,22 | 15,22 | 122 | 1967 |
| RA | F | 2,62 | 39,16 | 14,96 | 120 | 1894 | 38,91 | 14,86 | 120 | 1906 |
| AA | S | 11,94 | 268,32 | 22,46 | 90 | 1822 | 242,25 | 20,28 | 99 | 2018 |
| SA | S | 38,51 | 824,03 | 21,40 | 100 | 1943 | 737,50 | 19,15 | 111 | 2171 |
| SY | S | 14,54 | 311,95 | 21,46 | 104 | 2011 | 275,13 | 18,93 | 118 | 2280 |
| *M. kaiseri* |  |  |  |  |  |  |  |  |  |  |
| MA | F | 8,17 | 105,80 | 12,95 | 107 | 1736 | 95,71 | 11,72 | 119 | 1919 |
| RA | F | 7,06 | 88,95 | 12,60 | 111 | 1750 | 80,77 | 11,44 | 122 | 1927 |
| *M. ramanankasinae* |  |  |  |  |  |  |  |  |  |  |
| RA | F | 24,15 | 267,03 | 11,06 | 102 | 1614 | 216,84 | 8,98 | 126 | 1987 |
| *S. martini* |  |  |  |  |  |  |  |  |  |  |
| SA | S | 18,92 | 319,00 | 16,86 | 102 | 1857 | 235,23 | 12,43 | 129 | 2519 |
| *E. cf. insulicola* |  |  |  |  |  |  |  |  |  |  |
| RA | F | 30,23 | 319,07 | 10,55 | 141 | 2238 | 204,02 | 6,75 | 221 | 3500 |
| SA | S | 22,32 | 323,84 | 14,51 | 80 | 1565 | 181,23 | 8,12 | 143 | 2797 |
| *P. cf. vayssierei* |  |  |  |  |  |  |  |  |  |  |
| SA | S | 127,03 | 375,20 | 2,95 | 585 | 11411 | 578,17 | 4,55 | 380 | 7405 |

Size differences at emergence were analyzed for the 3 Baetidae species. We observed significantly larger size at emergence in deforested areas for *X. namorona* females (Kruskal-Wallis chi square = 8.2664, *p* = 0.0040, *n* = 39) and both sexes of *X. sartorii* (Kruskal-Wallis chi square = 5.1887, *p* = 0.0227, *n* = 41 for males, and chi square = 11.5474, *p* = 0.0007, *n* = 36 for females). No significant differences were observed for males of *X. namarona* (Kruskal-Wallis chi square = 1.9648, *p* = 0.161, *n* = 61) or for either sex of *D. merina* (Kruskal-Wallis chi square = 0.5016, *p* = 0.4788, *n* = 45 for males, and chi square = 0.1163, *p* = 0.7331, *n* = 48 for females).

### Secondary production

Mean biomass ranged from < 3 mg DW m^-2^ (*X. namarona* in a forest site) to almost 130 mg DW m^-2^ (*P. cf. vayssierei* in *savoka*). Biomass of *D. merina* and *X. namarona* was significantly greater in *savoka* compared to forest (p = 0.083 and p = 0.052 respectively), whereas *X. sartorii* biomass was similar (Fig. 3B). Production estimates obtained by the instantaneous growth method were 2-44% higher than those derived from the size-frequency method for all species, except for *P. cf. vayssierei* (−54%). Differences were always lower in forest sites when compared with *savoka*. All three Baetidae species exhibited higher production in *savoka* compared to forest sites, being between ca 3.5 (*Xyrodromeus* species) and 4.5 (*D. merina*) times more productive (Fig. 3C). Annual turnover (P/B) ranged from less than 3 (*P. cf. vayssierei*) to more than 22 (both *Xyrodromeus* species in savoka), most of the values falling in the range 12-21.

**Fig. 3:**
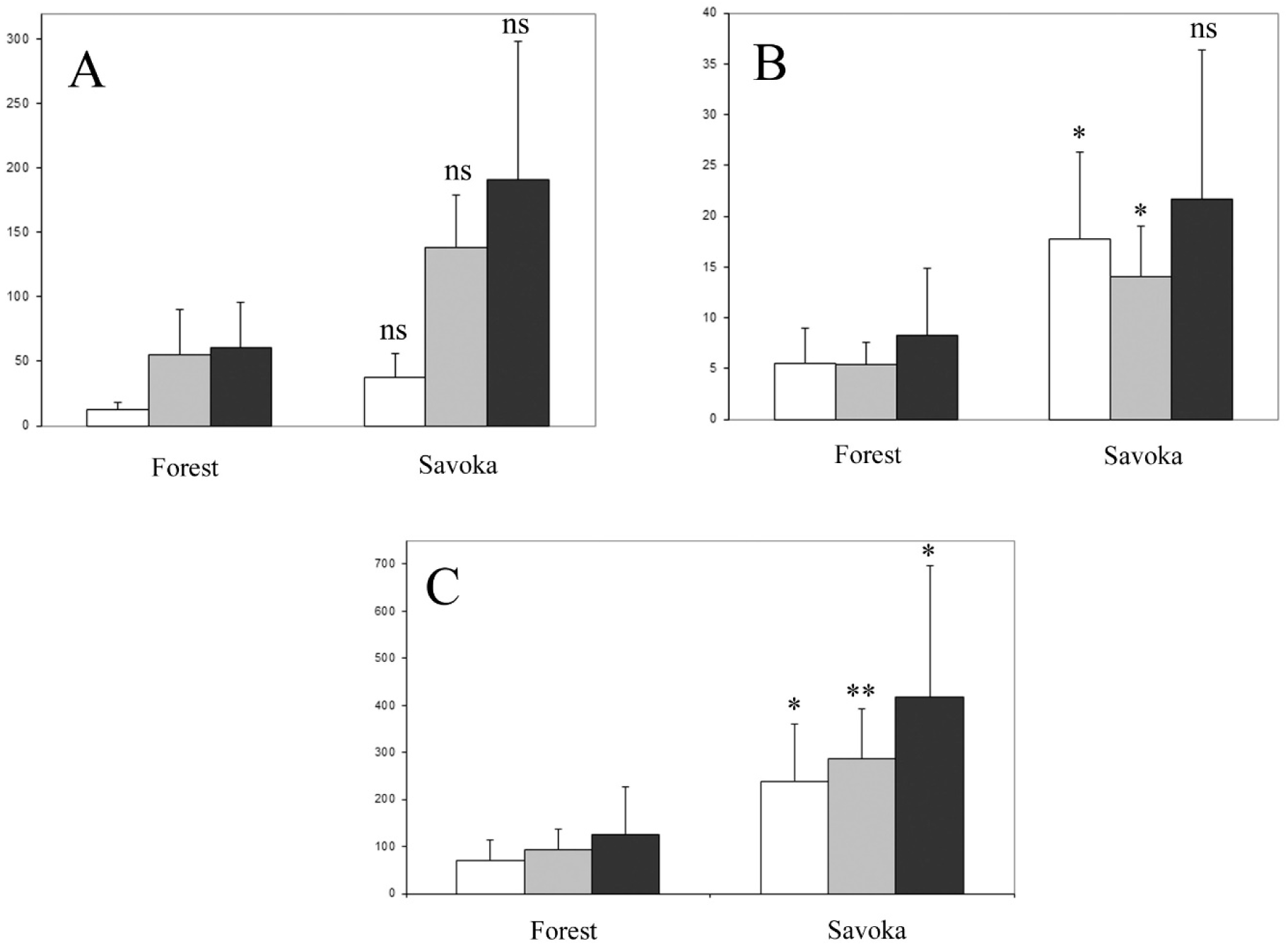
Comparison of results obtained for the 3 baetid species in forest (N=3) and in savoka (N=3). White bars: *D. merina*; grey bars: *X. namarona*; black bars: *X. sartorii*; values with SD; A: density (nb. ind. m^-2^), B: biomass (mg DW m^-2^), C: production (mg DW m^-2^ y^-1^) (instantaneous growth method); ns: non significant; * significant at p<0.05; ** significant at p<0.01.

## Discussion

The majority of studies reporting population-based secondary production come from freshwater and marine ecosystems (Benke 2010), but production estimates for tropical stream insects are scarce. These are the first we are aware of from the Afrotropical region, and the first quantitative data for any mayfly species belonging to Oligoneuriidae, Euthyplociidae, Tricorythidae or Teloganodidae. Although among the smallest species studied, the 3 Baetidae were the most productive in *savoka*, with average values (320-470 mg DW m^-2^ y^-1^) similar to those obtained for the larger *Elassoneuria* and *Proboscidoplocia*, the latter of which is among the largest mayflies on Earth. These values are, together with a *Pseudocloeon sp*. population on Papua New Guinea (Marchant and Yule 1996), among the highest reported values for Baetidae. Productivity of other species studied here (*M. kaiseri, M. ramanankasinae* and *S. martini*) were also high compared with other pannote mayflies such as *Leptohyphes* or *Tricorythodes* (Benke and Jacobi 1986, Jackson and Sweeney 1995, Ramirez and Pringle 1998), but lower than some *Caenis* in which production in subtropical areas can exceed 6 g DW m^-2^ y^-1^ (Yan and Li 2007).

Our results clearly indicate that mayfly communities in *savoka* are more productive than those in forest. This was mainly due to the increase in Baetidae biomass and production.

Among the most important factors typically influencing biomass and production are temperature and food availability (Benke 1984, 1993). The higher water temperatures measured in *savoka* streams (typically > 3°C) certainly play a role and are therefore of interest to studies of climate change (Winterbourn et al. 2008). A very interesting finding of this study is the fact that females of two dominant members of the community (*Xyrodromeus*) were larger at maturity in *savoka*. This was the opposite of what we expected (i.e. reduced size at emergence due to a faster development in warmer streams) based on earlier work (Sweeney and Vannote 1978). Our results suggest that *Xyrodromeus* species are better adapted to *savoka*, where environmental conditions are more suitable and where females are larger and presumably produce more eggs. This implies that colonisation of forest streams by *Xyrodromeus* could be secondary, with the reduced size at emergence and longer life cycle duration indicative of suboptimal conditions found in forest. Size of the third dominant Baetidae, *D. merina*, was not different in forested and deforested areas. It is larger than the two *Xyrodromeus* species, and its mouthparts are less specialized, suggesting it is more opportunistic and adaptive.

Our results indicate 4-5 times lower annual turnover ratios (P/B) than in tropical Asia, where Salas and Dudgeon (2003) found P/B values ranging from 77-109 for Baetidae in two Hong Kong streams. This difference was mainly due to much lower biomass and a cohort production interval ca two times shorter than in our study. These are the only production data published for tropical Baetidae, and this highlights again the scarcity of published data; we have no results on Baetidae P/B from the Neotropics, thus any comparison is impossible.

With the exception of *P. cf. vayssierei*, all studied species were multivoltine. *E. cf. insulicola* is the one which presents the most important difference, with a life cycle ca 2 times longer in forest compared to savoka. *P. cf. vayssierei* belongs to a tropical family (Euthyplociidae) which encompasses the largest mayflies on earth (Elouard et al. 2003), and few studies have been done of their life cycle and secondary production, with the exception of *Euthyplocia hecuba* which has a 2 years life cycle in Central America (Sweeney et al. 1995). Development time for the 3 baetid species was comparable to *Pseudocloeon sp*. on Bougainville Island, Papua New Guinea (Marchant and Yule 1996), but much longer than for *Acerpenna sp.* (23 days) in Costa Rica (Jackson and Sweeney 1995) or even *Baetis sp.* (41 days) in subtropical North America (Benke and Jacobi 1986). The two Tricorythidae species also present comparable life cycle interval compared to representatives of their sister family in the Neotropics, Leptohyphidae, such as *Leptohyphes sp*. or *Tricorythodes* sp. (Jackson and Sweeney 1995, Ramirez and Pringle 1998).

Some species were clearly associated with the forest *(Manohyphella, Madecassorythus)* and were rare in the *savoka*, whereas others *(Spinirythus, Proboscidoplocia)* showed the opposite pattern. In term of functional feeding groups, we can observe a shift from collector-gatherers to scrapers between forest and savoka streams (fig. 4), indicating that open streams offer greater food availability (algal periphyton) for the *Xyrodromeus* species. These findings are in agreement with the general pattern of shifting between allochtonous food resources dominant in forested areas to autochtonous food resources dominant in open canopy areas (Vannote et al. 1980). This is confirmed by the study of Bixby et al. (2009) who studied algal communities in Ranomafana area in Madagascar and showed that algal community in open-canopy streams, although less diverse, is more abundant than in forest ones, hence offering more opportunities to scrapers.

**Fig. 4:**
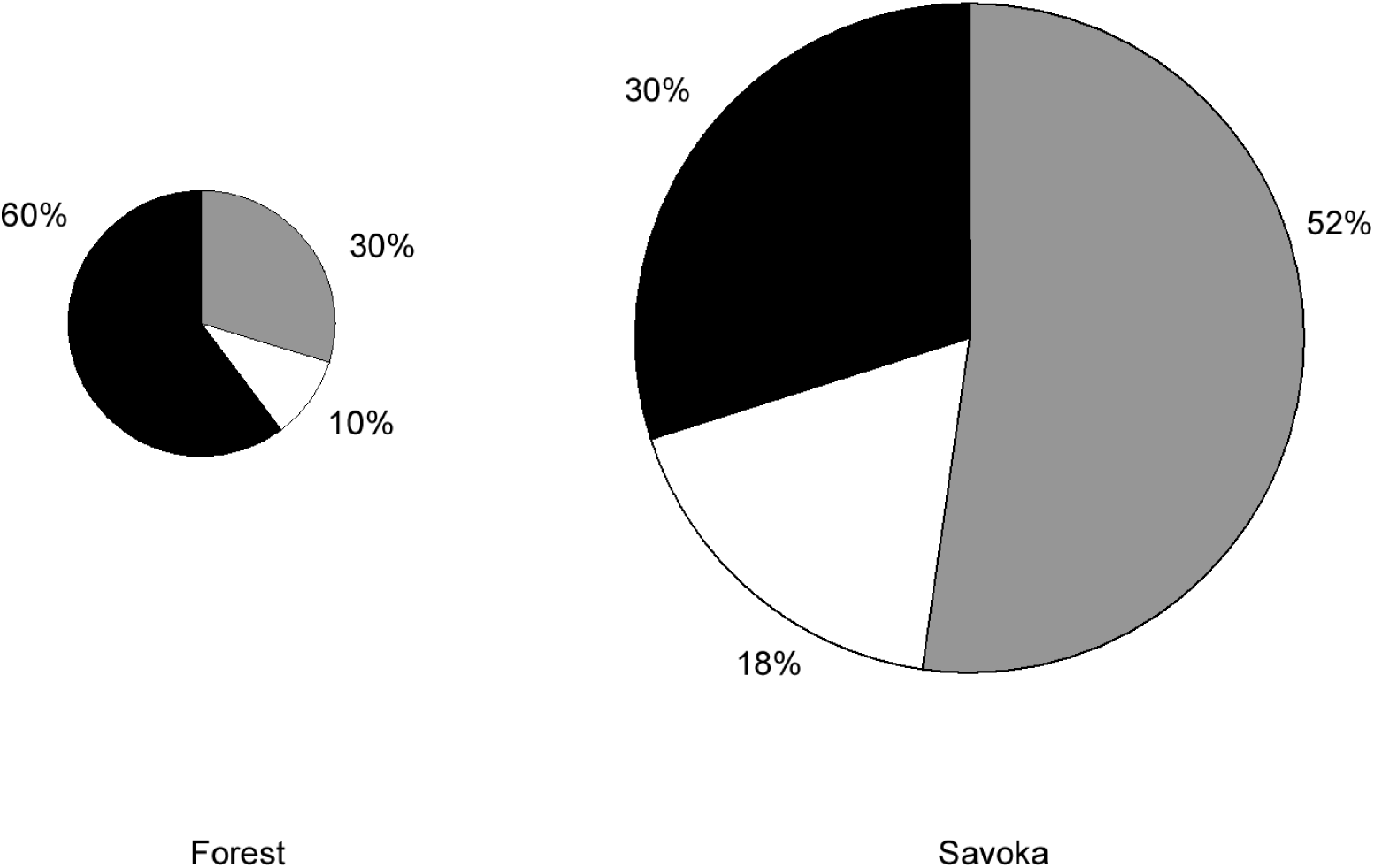
Mayfly production differences expressed by functional feeding groups between forest and savoka. Grey: scrapers, black: filterers, white: collector gatherers. Pie size proportional to mean production: forest = 971 mg DW m^-2^ y^-1^; savoka = 2134 mg DW m^-2^ y^-1^.

For species that occur in both habitats, mean densities of the three Baetidae species were higher in *savoka* than in forest although not significant. These findings are similar to those of Benstead et al. (2003), or Benstead and Pringle (2004), who found higher densities of Baetidae *(Afroptilum sp.)* and lower densities of *Elassoneuria sp*. in savoka than in forest in Madagascar. Their results from Tricorythidae are difficult to interpret because the single reported genus name (*Tricorythus)* probably encompasses 3 genera (*Madecassorythus* being common in forest and *Spinirythus* in *savoka* M. Sartori, unpubl. results). Results with standing biomass are broadly congruent with those obtained with densities, and the increased standing biomass of Baetidae in *savoka* was significant for the two baetid *D. merina* and *X. namarona*. But our values remain much lower than those from Benstead and Pringle (2004) with *Afroptilum sp*. although densities were comparable. The absence of the length-mass relationships they used for biomass calculation hinders us to discuss in more detail these discrepancies. In other words, scraper baetid nymphs are favoured in these disturbed environments when compared to original, forested ones. Similar results have been recently found in headwater streams in Costa Rica, where Baetidae are shown to be among the taxa favoured by deforestation (Lorion and Kennedy 2009). Knowing that the rate of deforestation throughout the world is still dramatic in the tropics, we can predict that this situation will certainly favour, to a certain point, Baetidae species compared to other mayflies, and that representatives of this family will be more and more dominant in tropical Ephemeroptera communities.

## Acknowledgments

Funding was provided by a PhD scholarship from the University of Lausanne to RO and the International Foundation for Science (Stockholm). We thank the *Association Nationale pour la Gestion des Aires Protégées* and the *Direction des Eaux et Forêts* for facilitating research in the Andasibe area. The *Département des Eaux et Forêts de l’Ecole Supérieure des Sciences Agronomiques* helped with laboratory work in Madagascar. In Lausanne we thank Sandra Knispel (field), Olivier Glaizot (analysis and discussion), Jean-Luc Gattolliat (Baetidae identification and discussion), and Geneviève L’Eplattenier (laboratory) for their help. Comments by Michael T. Monaghan (IGB, Berlin) greatly improved the manuscript.

